# Extracellular membrane vesicles – previously unrecognized components of *Staphylococcus aureus* biofilms

**DOI:** 10.64898/2026.04.07.717111

**Authors:** Jinger Lei, Misaki Foster, Emery Ng, Erin S Gloag, Xiaogang Wang

**Author notes:** Correspondence (X. Wang).

## Abstract

*Staphylococcus aureus* is a leading cause of biofilm-associated infections, in which communities of bacterial cells are encased in an extracellular matrix composed of polysaccharides, proteins, and extracellular DNA (eDNA) that protect bacteria from host immune defense and antibiotics. Despite their importance, the mechanisms by which matrix components are released from bacterial cells and incorporated into the biofilm matrix remain poorly understood. Using a drip-flow biofilm system, we showed that MVs were associated with the biofilm matrix formed by *S. aureus* clinical isolate MN8. Proteomic analysis of biofilm matrix proteins and purified MVs showed that biofilm-derived MVs carried cytoplasmic, membrane, and extracellular proteins that closely resembled the protein composition of the biofilm matrix but differed significantly from MVs produced by planktonic cultures. Biofilm-derived MVs carried significantly higher levels of DNA than MVs from planktonic cultures, and MV-associated DNA was resistant to DNase treatment. Although strain MN8 is known to form polysaccharide-dependent biofilms, exogenously added DNase or proteinase K significantly impaired biofilm formation and integrity. Notably, these inhibitory effects were reversed by the addition of biofilm-derived MVs, which significantly restored biofilm formation in enzyme-treated cultures. Together, these findings provide evidence that *S. aureus* MVs are generated within biofilms, and that these MVs serve as an important resource of matrix components and contribute to biofilm formation.

**Importance:** Extracellular membrane vesicles (MVs) are important mediators of intercellular communication and have been implicated in the physiology and pathogenesis of bacterial infections. While MV production in *S. aureus* planktonic cultures has been recognized for over one decade, their presence and function in *S. aureus* biofilm formation have remained unexplored. Here, we report for the first time the purification and characterization of MVs derived from *S. aureus* biofilms. Our studies demonstrate that *S. aureus* MVs are important components of the biofilm matrix that contribute to biofilm formation by serving as key carriers of matrix proteins and eDNA. This work advances our limited understanding of MVs in Gram-positive bacteria and reveal a previously unrecognized mechanism underlying *S. aureus* biofilm formation.

## Introduction

*Staphylococcus aureus* is a leading cause of biofilm-associated infections(1), including those related to medical implants and osteomyelitis, both of which present significant clinical challenges(2, 3). Biofilms are a structured community of microbial cells encased within a self-produced extracellular matrix which play a key role in chronic infections by protecting bacteria from both the immune system and antibiotics(4). The biofilm extracellular matrix (ECM) is comprised of proteins, polysaccharides, and extracellular DNA (eDNA). Although the production of Poly-N-acetyl-β-(1–6)-glucosamine (PNAG), also known as polysaccharide intercellular adhesion (PIA), is required for biofilm formation in some *S. aureus* strains, particularly methicillin-sensitive *S. aureus*, most methicillin-resistant *S. aureus* (MRSA) isolates rely primarily on proteins rather than polysaccharides for biofilm formation(5–8). Recent proteomic analyses of *S. aureus* biofilm ECMs have revealed that both protein- and polysaccharide-dependent biofilms share a core exoproteome enriched in cytoplasmic proteins(9–12), which contribute to biofilm formation by moonlighting as ECM components. These proteins, together with extracellular DNA (eDNA), help maintain biofilm integrity through a complex network of electrostatic interactions(13, 14). More recently, this model has been extended to include lipoproteins that function as anchor points between eDNA and the bacterial cell surface(15). Despite significant progress in characterizing ECM components, the mechanisms governing the release and incorporation of matrix components, and the forms in which they are present within biofilm matrix remain poorly understood.

Extracellular membrane vesicles (MVs) are nano-sized, spherical particles that are produced by eukaryotes, archaea, and bacteria(16). MVs have been recognized as important cargo carriers in *S. aureus* planktonic cultures(17–25), comprising cytoplasmic, membrane, and extracellular proteins, as well as glycopolymers and nucleic acids(17, 21–23, 26–28). MVs generated from *S. aureus* planktonic cultures induce cytotoxicity in various cell types(23, 29–31), elicit pro-inflammatory responses(21, 25, 31–34), and promote bacterial survival in human and animal blood(34). Moreover, *S. aureus* MVs have been detected during in vivo acute infection(35), shedding light on their physiological relevance in disease settings. Whereas these findings suggest a role of MVs in *S. aureus* virulence, whether MVs are produced in staphylococcal biofilms and contribute to biofilm biology remain largely unexplored.

Biofilms represent a sessile, matrix-embedded lifestyle underlying many chronic and device-related infections, where bacteria persist despite activated immune defense and antibiotic treatments. These distinct conditions may profoundly influence MV production, composition, and function. MVs produced within *Candida albicans* biofilms carry specific cargo proteins that contribute to the development of biofilms and their drug resistance(36–39). In addition, MVs from *C. albicans* biofilms cultivated for different time points exhibit temporal differences in inducing host cell responses(40). In Gram-negative bacteria, outer membrane vesicles (OMVs) have been characterized in biofilms of multiple bacterial species, such as *Pseudomonas aeruginosa*(41)*, Vibrio cholerae*(42)*, and Helicobacter pylori*(43), and these OMVs play a variety of roles in biofilm development(44). For instance, OMV production is induced by quinolone signal in *P. aeruginosa* biofilms, and these OMVs promote biofilm dispersal through encapsulation of enzymes capable of degrading matrix components(45). Conversely, OMVs generated within *V. cholerae* biofilms contribute to biofilm matrix assembly by carrying key matrix proteins and polysaccharide(42).

Although considerable progress has been made in understanding the role of MVs in biofilm formation across diverse microorganisms, their production, composition, and function in Gram-positive bacteria remain poorly characterized. *Bacillus subtilis* was shown to produce MVs in *in vitro* biofilms, but the cargo of these MVs and their role in biofilm formation remains unexplored(46). Two studies visualized MVs in electron micrographs of static *S. aureus* biofilms grown in tissue culture plates, but MVs were neither purified nor functionally characterized(47, 48). In this study, we provide the first characterization of MV production within *S. aureus* biofilms, utilizing a well-established drip-flow biofilm system. We provide evidence that *S. aureus* MVs are important carriers of key matrix components, including eDNA and proteins, and contribute to biofilm formation. Our study provides new insights into the roles of MVs in biofilm formation in Gram-positive bacteria.

## Results

### Purification of MVs from *S. aureus* biofilms using a drip-flow biofilm system

We used a drip-flow biofilm system(49–52) to cultivate *S. aureus* MN8 biofilms at 37°C. This system allows biofilm growth under low shear at the air-liquid interface, promoting high biomass. *S. aureus* MN8 biofilms formed on glass slides by 48 h (Fig. 1A), and biofilms stained with the DNA dye SYTO-9 were imaged with confocal laser scanning microscopy (CLSM) (Fig. 1B). To assess the presence of *S. aureus* MVs in biofilms, biofilms were harvested, and ultrathin sections of biofilms were examined by transmission electron microscopy (TEM). We observed MV release from *S. aureus* cell surface (Fig. 1C), and massive numbers of vesicles were present in the biofilm matrix (Fig. 1D).

**Figure 1.**
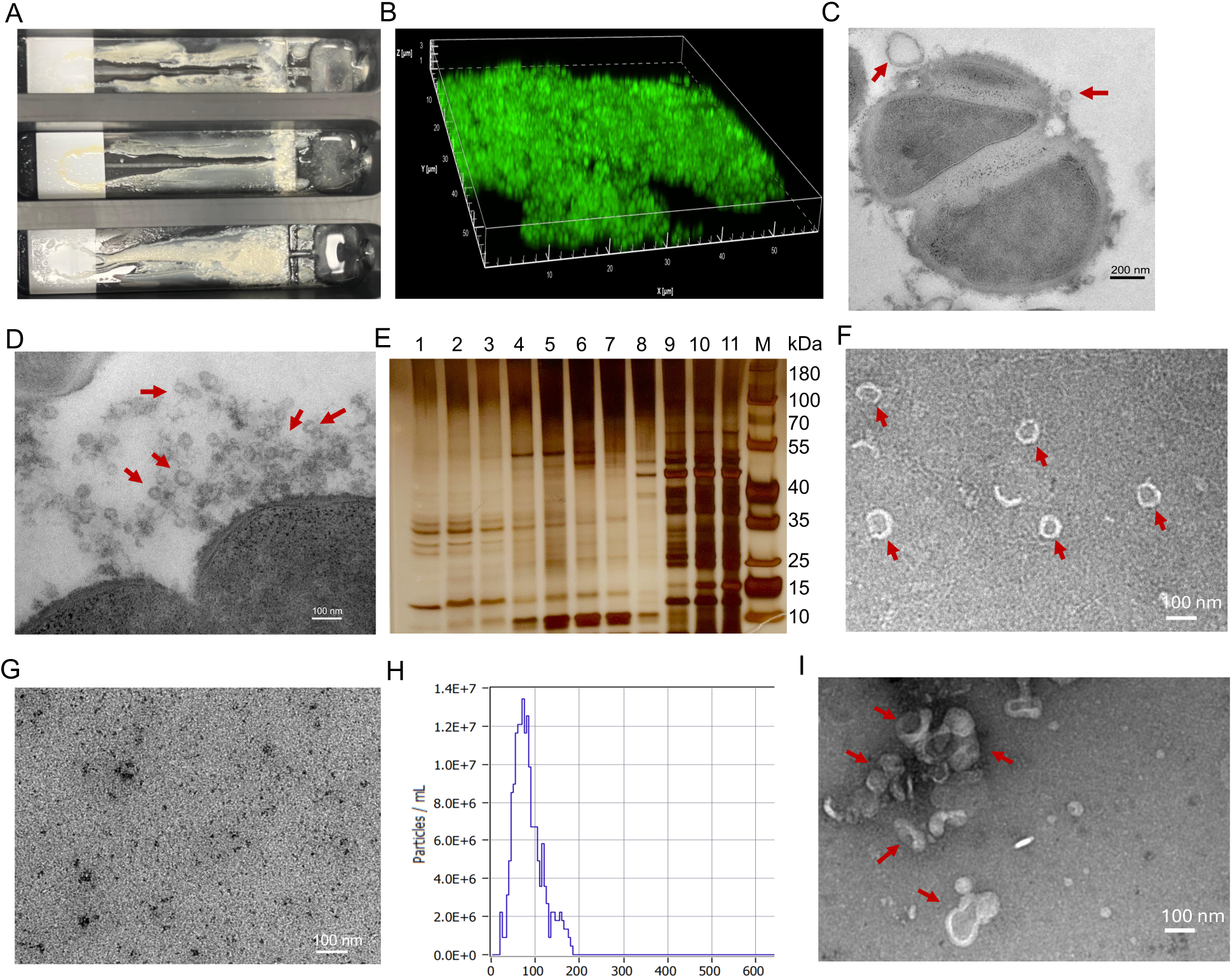
Generation of MVs in *S. aureus* MN8 drip-flow biofilms. **A***, S. aureus* MN8 biofilms formed on glass slides using a drip-flow bioreactor. **B**, Representative fluorescent image of MN8 drip-flow biofilms stained with SYTO-9 DNA dye. **C, D**, Ultrathin sections of MN8 drip-flow biofilms examined by TEM revealed MVs (indicated by red arrows) released from the cell (C) or present in the matrix (D). **E**, Silver staining of fractions recovered from the density gradient ultracentrifugation. **F-G**, TEM showed purified MVs (indicated by red arrows) in pooled fractions 2 to 7 (F), but not in fractions 8-11 (G). **H**, The size distribution of purified MVs was measured by nanoparticle tracking analyzer. **I**, MVs (indicated by red arrows) purified from effluent medium of drip-flow biofilms were imaged by TEM.

Crude MVs from the filter-sterilized supernatants of biofilm homogenates were pelleted by ultracentrifugation and subjected to OptiPrep density gradient ultracentrifugation to remove membrane fragments and protein aggregates. Consecutive Optiprep fractions (20 µl) were subjected to SDS-PAGE, and the proteins were silver-stained. OptiPrep Fractions with similar protein profiles (fractions 1-7 and 8-11; Fig. 1E) were pooled and diafiltrated in PBS. TEM imaging confirmed the presence of MVs in pooled fractions 1-7 (Fig. 1F), but not in fractions 8-11 (Fig. 1G). Nanoparticle Tracking Analysis (NTA) showed that purified MVs had an average size of 80 nm (Fig. H).

To determine whether MVs were also shed from the biofilm matrix during biofilm growth, the effluent medium from drip-flow biofilms was collected after 24 h and filter-sterilized. MVs pelleted by ultracentrifugation were further purified from OptiPrep density gradient ultracentrifugation. The recovery of MVs from effluent medium was confirmed by TEM imaging (Fig. 1I), indicating that MVs were also shed into the medium from *S. aureus* biofilms.

### Proteomic analysis of *S. aureus* MN8 biofilm matrix and biofilm-derived MVs

We analyzed protein contents of MN8 biofilm matrices, as well as MVs purified from MN8 drip-flow biofilms and planktonic cultures by using liquid chromatography-tandem mass spectrometry (LC-MS/MS). The *S. aureus* MN8 chromosome is 2,882,664 bp in size and comprises 2,833 predicted protein-coding sequences. This includes 1360 cytoplasmic proteins, 715 cytoplasmic membrane proteins, 40 cell wall-associated proteins, 98 extracellular proteins, and 619 proteins with unknown subcellular localization. We identified 747 proteins in the biofilm matrix and 730 proteins in MVs purified from biofilms (Table S1). Subcellular localization predictions indicated that most proteins in the biofilm matrix were classified as cytoplasmic, followed by membrane-associated, extracellular, and cell wall-associated proteins (Fig. 2A). Similarly, proteins identified in biofilm-derived MVs were predominantly cytoplasmic, followed by membrane-associated, extracellular, and cell wall-associated proteins (Fig. 2A). Of the 56 predicted lipoproteins in the MN8 genome, 31 were detected in the biofilm matrix and 35 were in biofilm-derived MVs (Table S2). 576 proteins were shared by biofilm matrix and biofilm-derived MVs (Table S3). Among shared proteins, 417 proteins (72.3%) were cytosolic, representing 74.4% and 89.3% of cytosolic proteins identified in the biofilm matrix and biofilm-derived MVs, respectively (Fig. 2B). 19.6% of shared proteins were membrane-associated, representing 83.6% of membrane proteins identified in biofilm matrix (Fig. 2B). 29 of 576 (5.02%) shared proteins were extracellular proteins, representing 83% and 85% of extracellular proteins present in biofilm matrix and biofilm-derived MVs, respectively (Fig. 2B). Many of these extracellular proteins are virulence factors, such as leukocidins LukAB and HlgBC, β-type phenol-soluble modulin (PSMβ1), chemotaxis inhibitory protein CHIPS, and proteases staphopain A and B.

**Figure 2.**
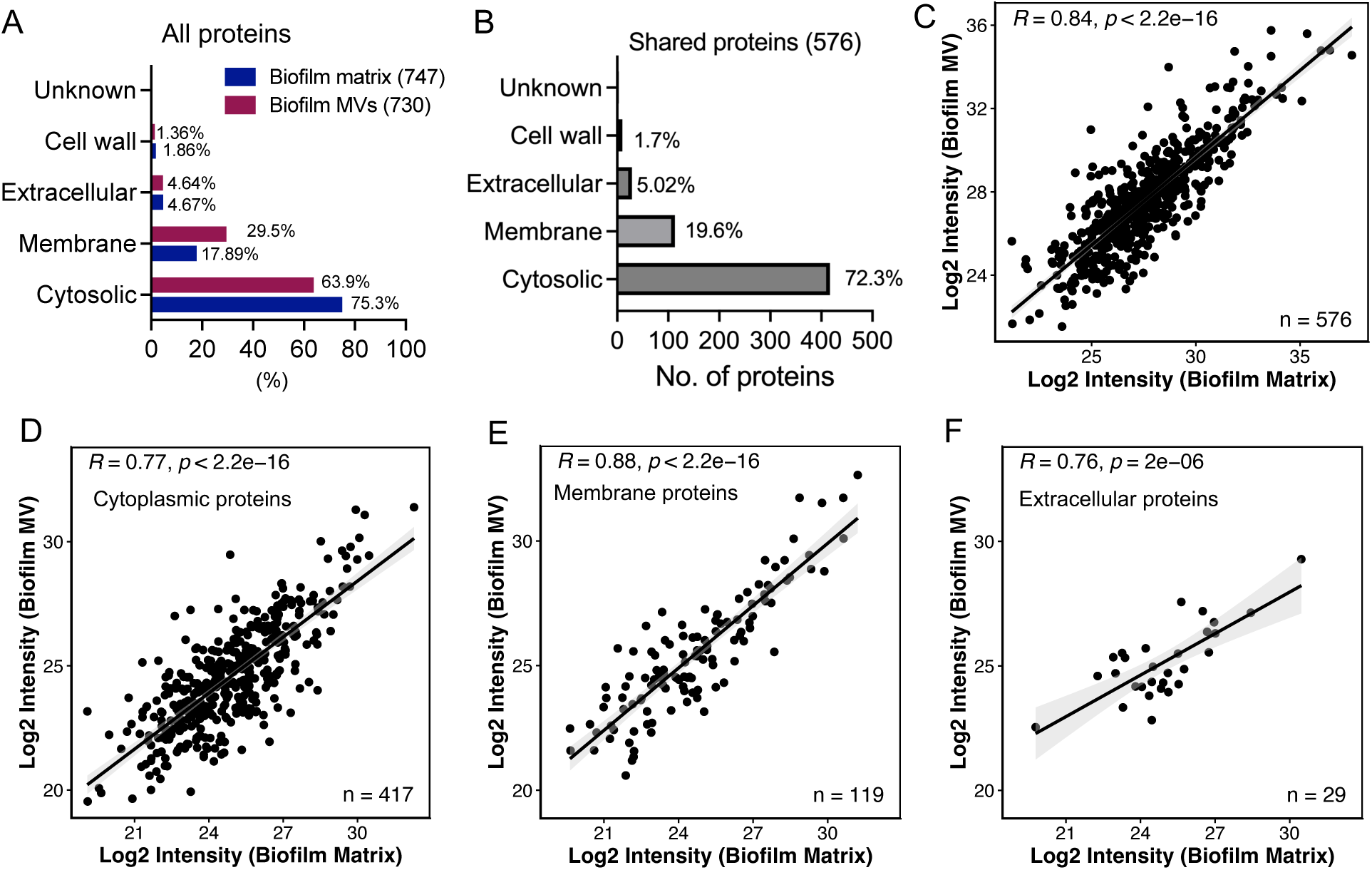
Proteomic analysis of MVs purified from *S. aureus* MN8 drip-flow biofilms and biofilm matrix. **A**, Subcellular localization of total proteins identified from purified MVs and biofilm matrix. **B**, Subcellular localization of proteins shared by purified MVs and biofilm matrix. **C-F**, Correlation of protein abundance between MVs and the biofilm matrix for proteins shared between two groups. Plots show correlations for (C) all shared proteins, (D) cytoplasmic proteins, (E) cell membrane proteins, and (F) extracellular proteins. Distributions are shown with linear regression lines.

Due to the high similarity between the protein contents of biofilm-derived MVs and the biofilm matrix, we hypothesized that the relative abundance of proteins in the biofilm matrix dictates their incorporation into biofilm-derived MVs. To test this hypothesis, we performed a correlation analysis of 576 shared proteins between the two groups, comparing the relative abundance of individual proteins in the biofilm matrix with their abundance in biofilm-derived MVs. Similar correlation coefficients were observed when all MV-associated proteins were analyzed together, as well as when cytoplasmic, membrane, and extracellular proteins were analyzed separately (Fig. 2C-2F).

### Comparison of MV proteomes from biofilm and planktonic cultures

Biofilms and planktonic cultures represent two distinct bacterial growth modes. We analyzed protein cargo of MVs purified from MN8 planktonic cultures and compared them with biofilm-derived MVs. In total, 568 proteins were identified in MVs purified from MN8 planktonic cultures (Table S1), compared with the 730 proteins identified in biofilm-derived MVs. Among the planktonic MV proteins, 48.1% were cytoplasmic, 45.3% were membrane-associated, 4.91% were extracellular, and 0.87% were cell wall-associated. 410 proteins were shared by MVs from both culture conditions, but 320 and 158 unique proteins were identified exclusively in biofilm- and planktonic-derived MVs, respectively (Fig. 3A). Among the 320 unique proteins identified in biofilm-derived MVs, 272 were cytosolic proteins; the remaining proteins were membrane (27), extracellular (10), and cell wall-associated proteins (8) (Fig. 3B). In contrast, 78 and 71 unique proteins present in MVs derived from planktonic cultures were cytoplasmic and membrane proteins, respectively (Fig. 3B). Notably, several proteins involved in the survival of *S. aureus* under stress conditions, including catalase, alkyl hydroperoxide reductase C, acetolactate synthase, thioredoxin, thioredoxin reductase, and thiol peroxidase were only identified in biofilm-derived MVs.

**Figure 3.**
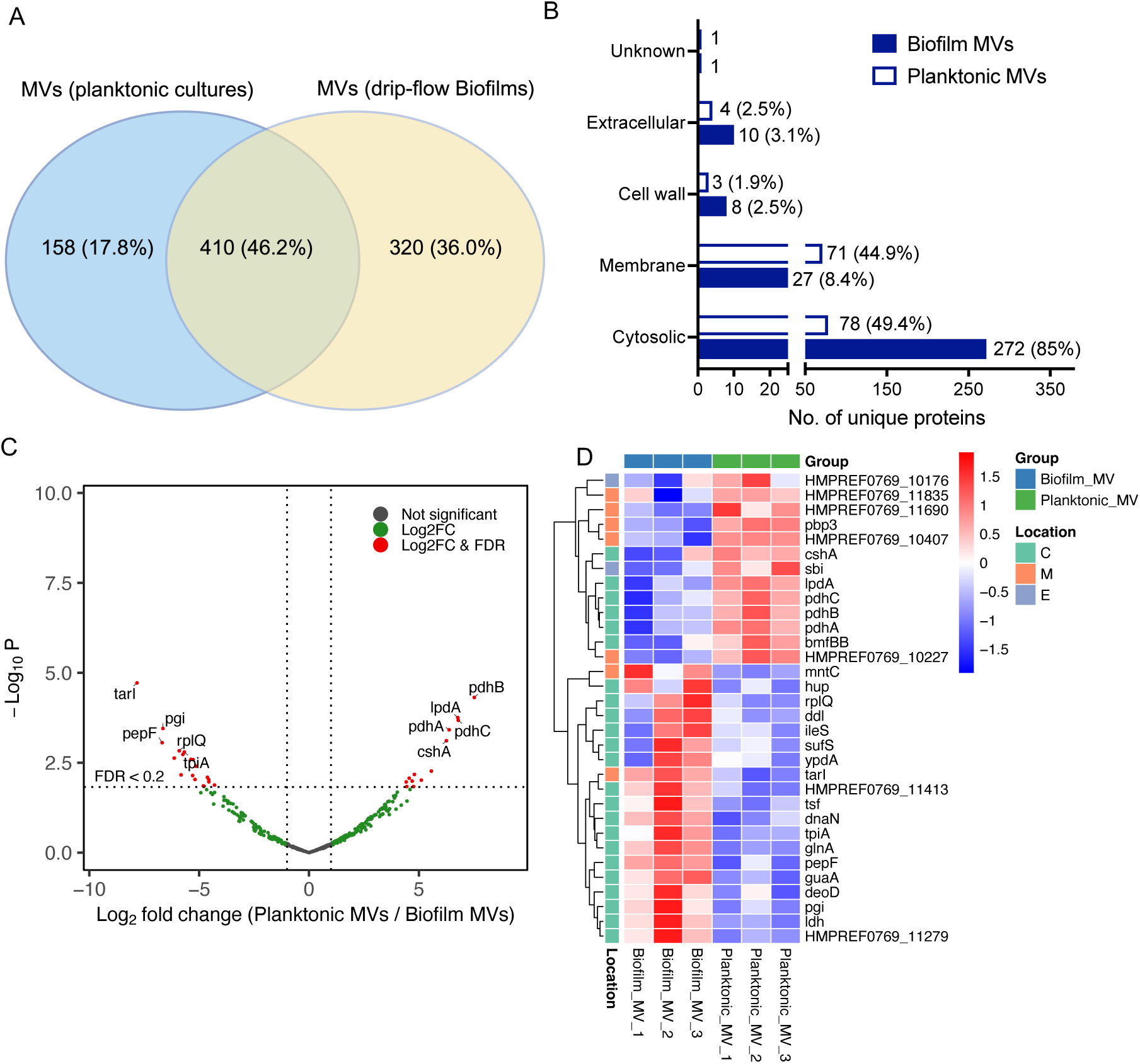
Comparison of proteome of MVs derived from *S. aureus* MN8 biofilms and planktonic cultures. **A**, Venn diagram illustrates the shared and unique proteins between the MVs isolated from drip-flow biofilms and planktonic cultures. The numbers within the diagram sections indicate the count and percentage of unique and shared proteins between two groups. **B**, Subcellular localization of unique proteins present in biofilm-derived MVs or MVs from planktonic cultures. **C**, Volcano plot shows the differential abundance of 410 shared proteins in MVs produced from biofilms and planktonic cultures. Each dot represents a protein, with the x-axis showing the log2 fold change (Log2FC) in protein intensities between biofilm-derived MVs and MVs from planktonic cultures. The y-axis shows the -log10 p-value of the change in intensity. Proteins with significant differential abundance at uncorrected *P* <0.02 and FDR <0.2 are highlighted in two colors. Ten of the proteins showing the greatest differential enrichment between biofilm-derived MVs or MVs from planktonic cultures are labeled by gene name. **D**, Heatmap view of 32 proteins with differential abundance (FDR <0.2) among the shared proteins between two groups. Each column indicates a biological replicate, and each row represents a protein. Predicted subcellular localizations are indicated in the legend. Proteins within each localization group were hierarchically clustered.

To further compare the shared proteins between two groups, we performed a volcano plot analysis using a fold change threshold of 2.0 (Fig. 3C). The analysis revealed 32 differentially abundant proteins showing significant differences (uncorrected *p<* 0.02, False Discovery rate (FDR)<0.2) between the biofilm-derived MVs and MVs from planktonic cultures. Among these, 19 proteins were more abundant in biofilm-derived MVs, while 13 had higher levels in the MVs from planktonic cultures (Fig. 3D). These analyses demonstrate that *S. aureus* MN8 biofilms produce MVs with a protein cargo that is compositionally distinct from those generated under planktonic conditions.

### Analysis of extracellular DNA associated with biofilm-derived MVs

We compared eDNA associated with biofilm-derived MVs versus MVs from planktonic cultures. Biofilm-derived MVs carried significantly higher levels of eDNA than MVs from planktonic cultures (Fig. 4A). We next measured eDNA present in MN8 biofilm homogenates before (S1) and after depletion of MVs (S2) by ultracentrifugation using a DNA-sensitive dye SYTOX Green(13). Depletion of MVs significantly reduced the fluorescence signal of SYTOX Green in the biofilm homogenates (Fig. 4B). In parallel, equal volumes of S1 and S2 fractions were subjected to agarose gel electrophoresis, and DNA was stained with SYBR safe DNA dye. Again, S2 fractions (MV-depleted) showed markedly lower DNA levels than S1 samples (Fig. 4C and 4D).

**Figure 4.**
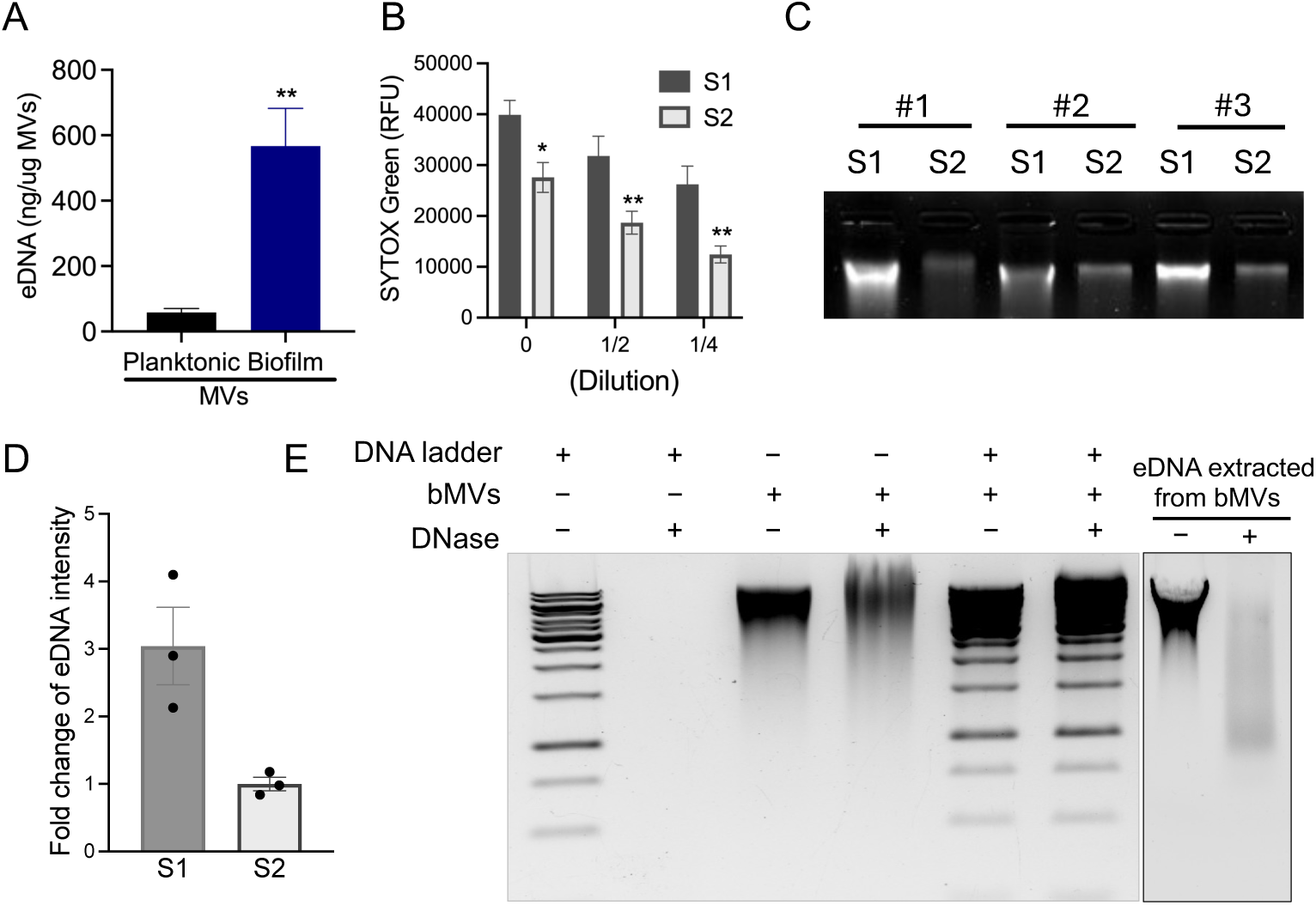
Association of eDNA with MVs derived from *S. aureus* MN8 biofilms. **A**, Quantification of eDNA associated with MN8 MVs purified from drip-flow biofilms or planktonic cultures. **B**, Quantification of eDNA present in filter-sterilized biofilm homogenates before (S1) and after (S2) MV depletion, measured by fluorescent DNA dye SYTOX Green. **C**, 30 μl aliquots of filter-sterilized biofilm homogenates before (S1) and after (S2) MV depletion were loaded onto DNA agarose gel and stained with SYBRO Safe DNA dye. Samples from three independent experiments were analyzed. **D,** Quantification of DNA signal intensities for S1 and S2 from triplicate samples. **E**, Biofilm-derived MVs (bMVs), 1 kb DNA ladder, or eDNA extracted from bMVs were treated for 1 h with or without 100 μg/ml DNase I, followed by agarose gel electrophoresis and staining with SYBR Safe DNA dye. Data are presented as mean ± standard error of the mean and analyzed using the Student’s *t* test. **, *P* < 0.01.

We treated biofilm-derived MVs for 1 h with 100 µg/ml DNase before analysis by DNA agarose gel. DNase treatment did not markedly reduce the levels of MV-associated eDNA. In contrast, the DNA ladder control and eDNA extracted from MVs were degraded by DNase I (Fig. 4E). Biofilm-derived MVs were incubated with 1 kb DNA ladder for 30 min before DNase I treatment. Notably, the exogenously added DNA ladder was protected from DNase I digestion following co-incubation with biofilm-derived MVs (Fig. 4E). Collectively, these data demonstrate that a substantial proportion of eDNA in *S. aureus* biofilm matrix is associated with MVs, and that MV-associated eDNA is protected from nuclease-mediated degradation.

### Detection of glycopolymers in MVs produced from *S. aureus* biofilms

Because PNAG is a major component of *S. aureus* MN8 biofilms, we examined the association of PNAG with MVs by immunodot blot using human antibodies against PNAG(53). PNAG was detected in WT MN8 cell lysates, but not in either biofilm-derived MVs or MN8Δ*ica* cell lysates (Fig. 5A), which served as a negative control. *S. aureus* MN8 produces type 8 capsular polysaccharide (CP8). We found that CP8 was associated with MVs derived from both biofilm and planktonic cultures (Fig. 5A). As controls, CP8 was present in WT MN8 cell lysates but absent in the capsule mutant (Δ*capHIJK)* lysates (Fig. 5A).

**Figure 5.**
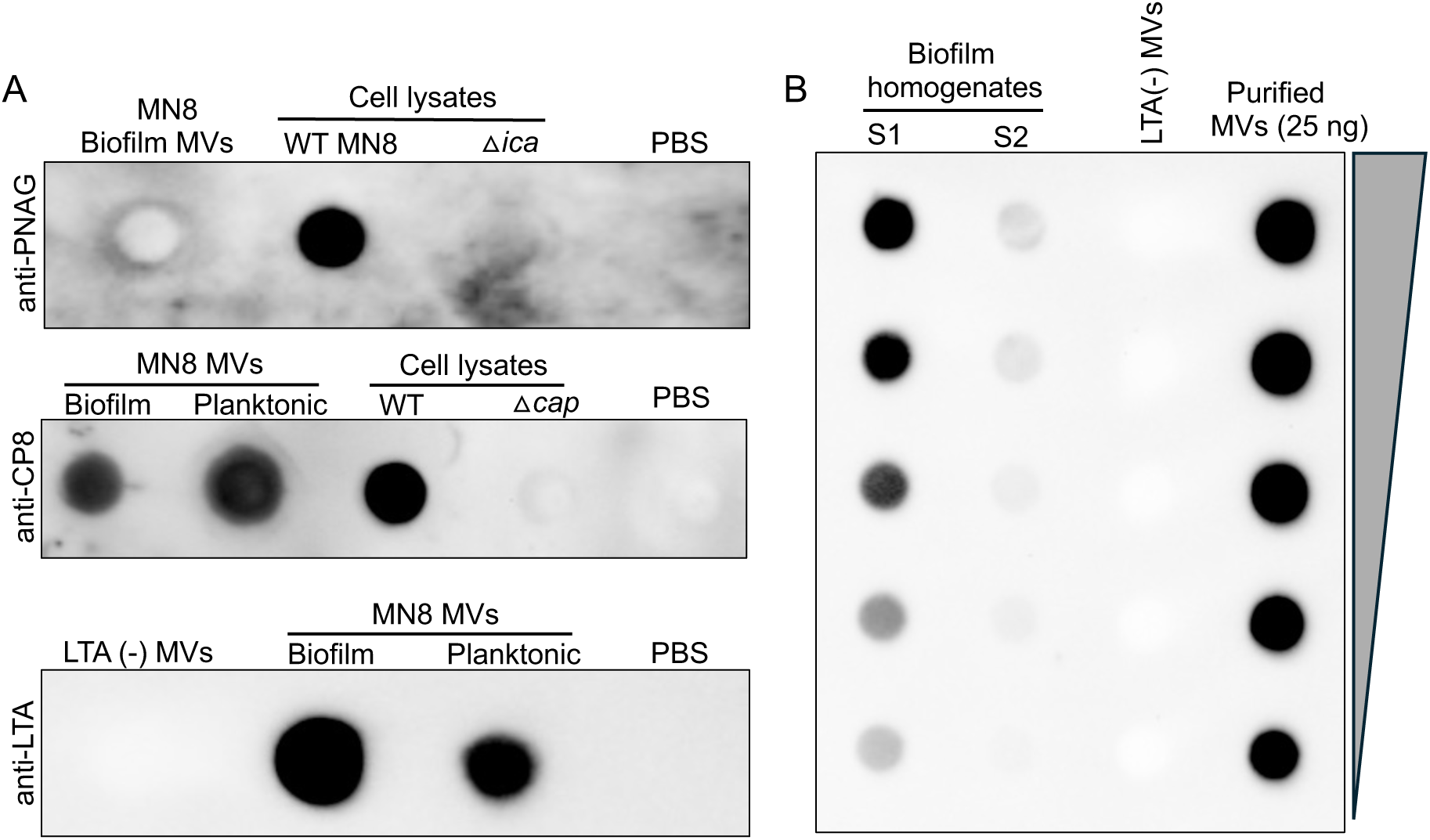
Detection of glycopolymers associated with *S. aureus* MN8 MVs. **A**, The presence of PNAG, CP8, or LTA in biofilm-derived MVs or MVs purified from planktonic cultures were determined by dot blots using specific antibodies. Where indicated, cell lysates from WT MN8 or relevant mutants deficient in PNAG or CP8 production were used as positive or negative controls. For LTA, MVs produced from an LTA-deficient *S. aureus* strain served as a control. PBS was included as a loading control. **B**, A total of 100 μl of two-fold serial dilutions of filter-sterilized biofilm homogenates or MVs (25 ng) purified from biofilms were applied to nitrocellulose membranes, and LTA was detected using anti-LTA antibodies. Serial dilutions of LTA-negative (–) MVs (2.5 µg) were used as a negative control. All experiments were repeated at least twice with similar results.

Lipoteichoic acid (LTA) is a membrane-anchored glycopolymer that is highly abundant in MVs produced from planktonic cultures(17). LTA was detected in as little as 25 ng MVs purified from either biofilms or planktonic cultures (Fig. 5A). In contrast, no signal was observed with up to 2.5 μg MVs produced from an LTA-deficient *S. aureus*(54), confirming the specificity of the assay. To determine whether LTA in the biofilm matrix was MV-associated, we depleted MVs from biofilm homogenates by ultracentrifugation. Dot blot analysis of 2-fold diluted samples showed the presence of LTA in original biofilm homogenates and purified MVs, but not in MV-depleted homogenates (Fig. 5B). Together, these results indicate that LTA molecules present in MN8 biofilms are predominantly associated with MVs.

### Assess the functional role of MVs in biofilm formation

Recent studies have suggested that, both eDNA and proteins remain important for biofilm formation in PNAG-positive *S. aureus* strains(10, 14, 15, 55). Strain MN8 is well known for its PNAG-dependent biofilms, and that eDNA is released during biofilm formation(55). To evaluate the contribution of these components on biofilm formation, we cultivated WT MN8 and its PNAG-negative strain Δ*ica* for 24 h in tissue culture plates containing TSB supplemented with 0.2% glucose and 1% NaCl. The resulting biofilms were then incubated for an additional 24 h in fresh medium containing either 20 µg/mL proteinase K or 100 µg/mL DNase I. Biofilm formation was quantified using crystal violet staining. As expected, MN8Δ*ica* mutant formed substantially less biofilm than the WT strain, confirming the role of PNAG in MN8 biofilm formation. Of note, both DNase I and proteinase K treatments resulted in significant reductions in biofilm biomass in both WT MN8 and the Δ*ica* mutant (Fig. 6A and 6B). Overall, these results demonstrate that PNAG, eDNA, and proteins are all key components of MN8 biofilms and contribute to its structural integrity.

**Figure 6.**
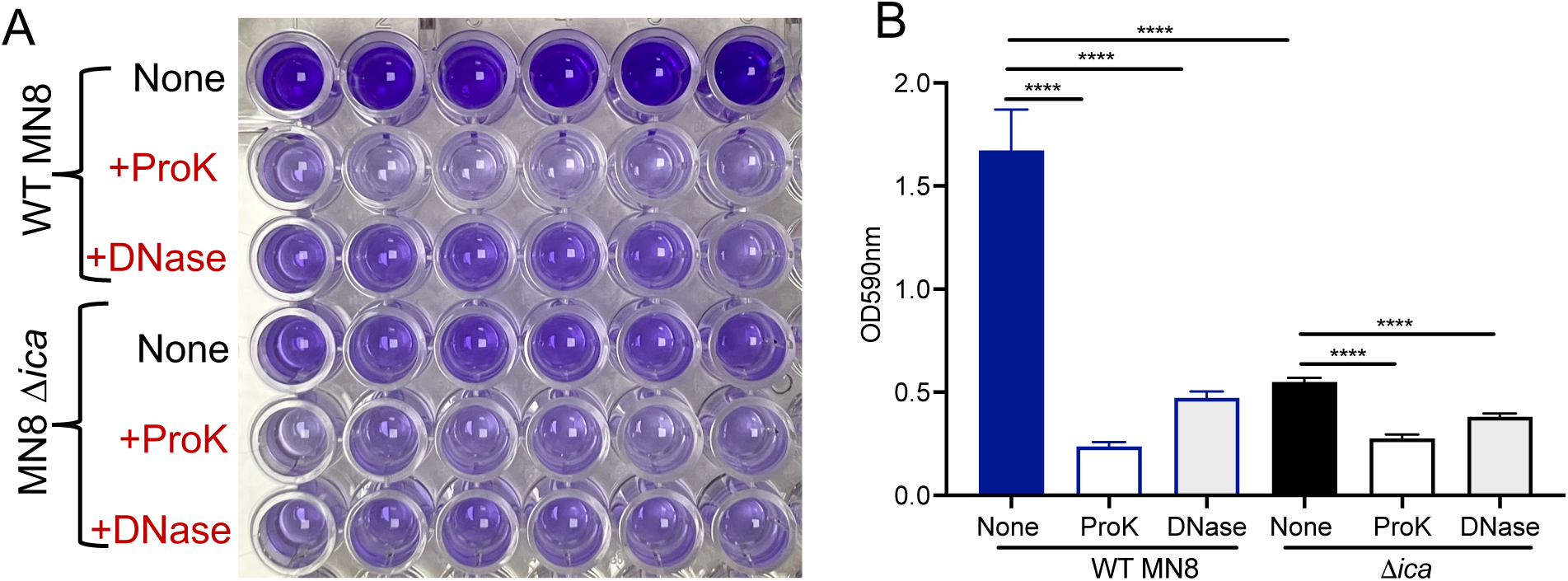
Proteins and eDNA are required for *S. aureus* MN8 biofilm formation. **A**, *S. aureus* MN8 and its Δ*ica* mutant biofilms were grown for 24 h in TSB supplemented with 0.2% glucose and 1% NaCl, followed by incubation for an additional 24 h in fresh medium with or without DNase I (100 µg/mL) or proteinase K (20 µg/mL). Biofilms were washed three times with PBS prior to staining with 0.2% crystal violet solution. **B**, Biofilm biomass was quantified by measuring the absorbance of the stained biofilms at 590 nm. Data are presented as mean ± SEM, and statistical analyses were performed using one-way ANOVA with Dunnett’s multiple comparison test. \*\*\*\**P* < 0.0001

Because MVs carry a substantial amount of eDNA and proteins present in MN8 biofilms, we evaluated whether MVs contribute to biofilm formation. 10 µg/mL of MVs purified from biofilms or planktonic cultures were added to 24-h MN8 static biofilms that had been pretreated with DNase I or proteinase K, followed by an additional 24 h incubation. Biofilm formation was then assessed by confocal microscopy. Addition of biofilm-derived MVs, but not MVs from planktonic cultures or PBS, significantly restored biofilm formation in static cultures pretreated with DNase I or proteinase K, as evidenced by increased biofilm biomass and biofilm thickness (Fig. 7A and 7B), Together, these results suggest that eDNA and proteins associated with biofilm-derived MVs serve as important structural components of *S. aureus* biofilms and contribute to biofilm formation.

**Figure 7.**
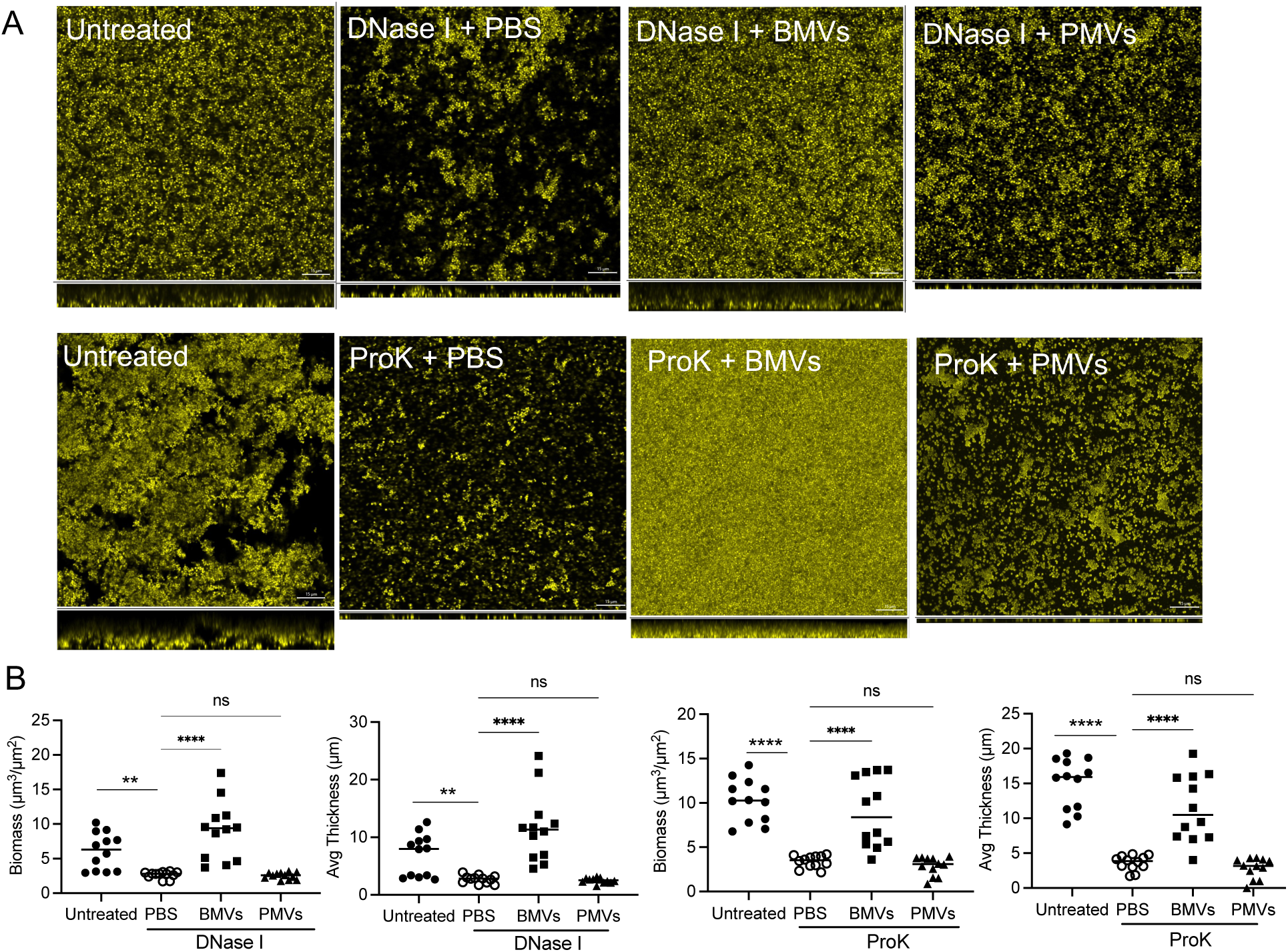
Biofilm-derived MVs promote biofilm formation. **A-B**, 10 μg/ml of MVs purified from drip-flow biofilms or planktonic cultures were added to 24-h MN8 biofilms that were pretreated for 6 h with proteinase K (20 μg/ml) or DNase I (100 μg/ml), followed by another 24 h incubation. Biofilms of each treatment were imaged by confocal microscopy (A), and biofilm thickness and biomass from 12 images of three biological replicates were quantified (B). Data are presented as mean ± SEM, and statistical analyses were performed using one-way ANOVA with Tukey’s multiple comparisons test. \**P* < 0.05, \*\**P* < 0.01, \*\*\**P* < 0.001, \*\*\*\**P* < 0.0001

## Discussion

Recent studies have shown that MVs produced by both Gram-negative bacteria and fungi play important roles in biofilm development, including matrix assembly, biofilm maturation, and modulation of host responses(36, 38–42, 45). In contrast, the generation and function of MVs within biofilms formed by Gram-positive bacteria, such as *S. aureus,* remain largely unexplored. We provide the first characterization of MVs derived from *S. aureus* biofilms. Our results indicate that *S. aureus* MVs produced during biofilm formation are not merely trapped in biofilms, but instead function as active contributors to biofilm formation. This work advances our limited understanding of the role that MVs play in biofilm formation in Gram-positive bacteria.

MN8 is known to form PNAG-dependent biofilms, as disruption of the *icaADBC* operon responsible for PNAG synthesis impairs the biofilm formation(56). We showed that proteinase K and DNase I treatments also significantly impaired MN8 biofilm formation, indicating that both proteins and eDNA are also important structural components of MN8 biofilms. Recent reports indicate that both eDNA and proteins are present within PNAG-dependent biofilms, and that cytoplasmic proteins have been implicated in moonlighting functions within biofilm matrix by interacting with negatively charged eDNA to contribute to biofilm integrity(10, 14, 55, 57, 58). Our proteomic analysis revealed 562 cytoplasmic proteins in MN8 biofilm matrix. Many previously identified moonlighting proteins(10) were detected, such as glyceraldehyde-3-phosphate dehydrogenase, enolase, transketolase, and lactate dehydrogenase. In addition, we identified 131 membrane-associated proteins, including 31 lipoproteins, such as MntC, FhuD2, PrsA, and SaeP. Although the contribution of lipoproteins to *S. aureus* biofilms remains ill-defined, lipoproteins mediate TLR2-mediated immune activation. *S. aureus* biofilm infections trigger host inflammatory responses(59–62). Moreover, several lipoproteins identified in this work, including PrsA and SaeP, were previously shown to bind eDNA within the *S. aureus* LAC biofilm matrix(15), suggesting a dual role in biofilm structural integrity and host immune activation. Collectively, our findings provide new evidence that incorporation of cytoplasmic proteins and lipoproteins into the extracellular matrix also occurs in PNAG-dependent biofilms, suggesting a conserved strategy used by *S. aureus* to assemble the biofilm matrix.

We previously demonstrated that *S. aureus* MVs produced in planktonic cultures serve as a secretory system enabling bacteria to transport their protected cargo to the environment and host cells(17, 21, 23, 63). Whether these findings extend to MVs generated within biofilms is unclear, as the bacterial cells cultivated under the two conditions differ substantially in physiology, composition, and immune dynamics. Notably, 77% of proteins present in the MN8 biofilm matrix were identified in biofilm-associated MVs, and the intensity-based correlation analysis also revealed a strong positive correlation between proteins detected in biofilm-derived MVs and biofilm matrix. Of the 562 cytoplasmic proteins identified in the MN8 biofilm matrix, 418 were enriched in biofilm-derived MVs, accounting for 89.3% of cytoplasmic proteins identified in biofilm-derived MVs. Over 80% of membrane and extracellular proteins identified in MN8 biofilm matrix were also present in biofilm-derived MVs, including lipoproteins and toxins, such as LukAB, HlgCB, PSMβ1, and TSST1. These results suggest that MVs contribute substantially to the protein composition of the *S. aureus* biofilm matrix and may also play a role in biofilm-associated infection.

Striking differences were observed in the MV proteome composition between biofilm and planktonic conditions. Biofilm-derived MVs carry 162 more proteins than MVs from planktonic cultures. 63.9% of proteins identified in biofilm MVs were cytoplasmic, and 29.5% of the proteins were membrane-associated. In contrast, the proteome of MVs purified from MN8 planktonic cultures was comprised of 48.07% cytoplasmic and 45.26% membrane-associated proteins. Bacterial biofilms represent a distinct lifestyle from planktonic cells, as evidenced by their different gene expression profiles(64). The differences in protein expression under the two growth conditions likely contribute to the observed variation in MV protein cargo. Moreover, proteins, such as catalase(65, 66), alkyl hydroperoxide reductase(66), thioredoxin, and flavohaemoglobin(67) contribute to biofilm formation under stress conditions, and these proteins were not detectable in MVs from planktonic cultures. Because MV production by planktonic *S. aureus* is also induced by certain environmental stress, such as oxidative stress, nutrient limitation, and antibiotic treatments(18), It is tempting to speculate that stress-related proteins associated with MVs play a role in bacterial adaptation.

As a structural component, eDNA interacts with proteins localized within the extracellular matrix and on the bacterial surface via electrostatic interactions(57, 68) to maintain the biofilm integrity and structure. Although recent studies have shown that several genes, including *gdpP*, *xdrA, sarA, and apt,* were involved in the release of eDNA within *S. aureus* biofilms(13, 55), the mechanisms by which eDNA is released remains poorly understood. We found that biofilm-derived MVs carry >10-fold more DNA than MVs from planktonic cultures, and depletion of MVs from biofilm homogenates resulted in >50% reduction in eDNA levels, indicating that a substantial proportion of eDNA within the biofilm matrix is associated with MVs. The majority of MV-associated eDNA was protected from DNase degradation, and that this protective effect was also applicable to exogenous DNA when it was pre-incubated with biofilm-derived MVs. These results suggest that eDNA may bind to the MV surface, and that this binding protects it from nuclease-mediated degradation. Kavanaugh et al. identified >60 eDNA-binding proteins, including cytoplasmic, membrane-associated, and extracellular proteins, in *S. aureus* biofilms and spent medium(15). Many of these proteins, such as Eap, Alt, MntC, PrsA, SaeP, Geh, Ltas, and the PSMβ1 peptide, were also enriched in biofilm-derived MVs. Thus, eDNA may be protected from nucleases through interactions with these DNA-binding proteins associated with MVs.

Bacterial biofilm formation is a dynamic process, and biofilm dispersal is a tightly regulated. Degradation and disruption of the extracellular matrix are essential elements for the dissemination of infection(69). *S. aureus* PSM peptides play a critical role in biofilm structuring and detachment(70). In addition, enzymes such as proteases and nucleases have been implicated in extracellular matrix degradation and dispersal(68, 71–73). However, the spatial and temporal regulation of these enzymes during biofilm maturation remains poorly understood.

In this study, we purified MVs from drip-flow biofilms cultivated for 48 h, a time point that may precede the dispersal stage. Periasamy et al. used a flow cell system to show that *S. aureus* biofilm dispersal occurred after 72 h(70). Nonetheless, we detected a relative low abundance of proteases (staphopains A and B), thermonucleases (Nuc1 and Nuc2), and PSMβ1 peptides in biofilm MVs. The expression of dispersal factors, including proteases, nucleases, and PSMs, is positively regulated by the Agr quorum-sensing system(70), which becomes highly active during late-stage biofilm development. MVs may contribute to biofilm matrix assembly at early stages by delivering key components, including eDNA and proteins. During later stages, matrix-degrading enzymes may become enriched within MVs. Because high concentrations of PSMs can lyse MVs(74), PSMs produced in mature biofilms may disrupt MV structures, facilitating the release of matrix-degrading enzymes and promoting biofilm dispersal. Interestingly, in *Candida albicans* biofilms MVs play multiple roles in biofilm development, such that MV functions are mediated by specific cargo proteins(36). Further studies are needed to elucidate how MV biogenesis and cargo are regulated throughout *S. aureus* biofilm development and how differentially abundant groups of proteins in MVs influence biofilm development.

## Materials and Methods

### Bacterial strains and growth conditions

*S. aureus* strains (listed in Table S4) were cultivated with aeration in tryptic soy broth (TSB, Difco) or agar plates at 37°C. To grow biofilms, overnight cultures were sub-cultured to fresh TSB at a 1:1000 dilution and grown to an OD650 of 0.6. Bacterial cells were pelleted by centrifugation at 4000 x *g* and washed with sterile PBS. The cell pellet was resuspended and further diluted 1:500 in TSB supplemented with 0.2% glucose (TSBG) and inoculated into each chamber of the drip-flow system (BioSurface Technologies Corp), which can hold up to 6 glass slides with ∼125 cm^2^ of surface area^2^. After overnight incubation at 37°C to allow bacterial attachment, the planktonic cells from each chamber were removed from the effluent port, and 10% TSB supplemented with 0.2% glucose was continuously supplied via a peristaltic pump at a flow rate of 0.25 ml/min for 48 hours.

### Purification of *S. aureus* MVs from planktonic cultures and drip-flow biofilms

*S. aureus* MN8 was grown in TSB at 37°C to the post-exponential phase (OD_650nm_ = 1.2) phase of growth, and MVs from these planktonic cultures were purified as we described previously(17, 21, 75). The protein concentrations of purified MVs were determined with Bio-Rad protein dye. MV particles were quantified using a Zetaview Nanoparticle Tracking Analyzer (Particle Metrix) with the following settings: camera sensitivity, 80.0; shutter, 90.0; frame rate, 30 frames per second; temperature, 25°C. The analyses were performed with the in-built Zetaview software 8.02.31.

To purify MVs from *S. aureus* biofilms, the drip-flow biofilms were harvested in sterile PBS at 48 h and disrupted by homogenization. The homogenates were centrifuged at 8000 x g at 4°C for 30 minutes to pellet bacterial cells, and the resulting supernatants were filter-sterilized with a 0.45 um filter. MVs derived from biofilms were pelleted by ultracentrifugation of supernatants at 150,000 x *g* at 4°C for 3 h, followed by density-gradient ultracentrifugation to remove protein aggregates and membrane fragments. Fractions with a similar protein profile on SDS-PAGE were pooled, and the Optiprep medium was removed by diafiltration with PBS using a Centrifugal Filter Unit (10 kDa, polyether sulfone, Thermo Fisher). Purified biofilm MVs were filter sterilized, and their protein yield and particle numbers were measured by protein assay and NTA, respectively.

### Proteomic analysis of MVs and biofilm matrix

The supernatants of biofilm homogenates were prepared as described above. After concentration with an Amicon® Ultra Centrifugal Filter with 3 kDa MWCO (MilliporeSigma), the protein content was determined. 20 µg of biofilm supernatants, biofilm-derived MVs, or MVs purified from planktonic cultures were loaded onto 10% SDS-PAGE gels and stained with Coomassie blue. Gel sections containing proteins bands of each sample were recovered and subjected to a modified in-gel trypsin digestion procedure and analyzed by LC-MS/MS using electrospray ionization and an LTQ Orbitrap Velos Pro ion-trap mass spectrometer (Thermo Fisher Scientific). Peptides were detected, isolated, and fragmented to produce a tandem mass spectrum of specific fragment ions for each peptide. Three biological replicates of each sample were analyzed by LC-MS/MS. Peptide sequences were determined by matching the NCBIProt database of the *S. aureus* MN8 genome with the acquired fragmentation pattern by the software program SEQUEST (Thermo Fisher Scientific). To remove contaminants and reverse hits, data were filtered to maintain an average 0.8% and 0.1% false discovery rate for protein and peptide identifications, respectively. Protein intensity was quantified as the summation of intensity values (peak height intensity) of individual peptides matched to a given protein. For enhanced confidence in protein identification, individual proteins in each dataset were further filtered to maintain a minimum requirement of two total peptides and one unique peptide in at least two of the three biological replicates. Subcellular localization of the proteins was predicted by PSORTb v3.0.3.

Bioinformatic analyses were performed using the R package (v4.1.3). Protein intensities were log_2_-transformed to achieve an approximately normal distribution of the data. Correlations in relative protein abundance between biofilm matrix and biofilm-derived MVs were evaluated using Spearman’s rank correlation coefficient (r). Differential abundance analysis was performed for proteins shared between biofilm-derived MVs and MVs from planktonic cultures using R package limma on log2-transformed intensities, with missing values retained as N/A. Proteins that were only identified in one group were reported separately and were excluded from this analysis. P values were adjusted for multiple comparisons using the Benjamini–Hochberg method to control the false discovery rate (FDR). Proteins were considered significantly differentially abundant if they met an FDR < 0.2 (uncorrected *p*< 0.015). Volcano plots were generated by plotting the log2 fold change against the -log10 (adjusted p-value). Heatmap of differentially abundant proteins were generated to visualize abundance patterns using pheatmap in R. Missing values were imputed only for visualization purpose.

### Transmission electron microscopy

Purified *S. aureus* MVs were visualized by TEM as described (23). For TEM imaging of samples with fixation, MV samples adsorbed to carbon-coated grids were fixed in 1% (w/v) glutaraldehyde in PBS for 5 min before staining with 1% uranyl acetate.

For imaging *S. aureus* biofilm ultrathin sections, the drip-flow biofilm matrix was pelleted by centrifugation at 4000 x g, and the pellet was fixed for overnight at 4°C with 2.5% Glutaraldehyde/2.5% Paraformaldehyde in 0.1 M sodium cacodylate buffer (pH 7.4) and postfixed with 1% osmium tetroxide/1.5% potassium ferrocyanide. After incubation in 1% uranyl acetate, samples were sequentially dehydrated in ethanol (50%, 70%, and 90%) before they were soaked in propylene oxide for 1 h and infiltrated overnight with a 1:1 mixture of propylene oxide and Spurr’s resin. The following day the samples were embedded in Spurr’s resin and polymerized at 60°C for 48 h. Ultrathin sections were stained with lead citrate and examined in a JEOL 1200EX TEM. Images were recorded with an AMT 2k CCD camera.

### Immunodot blot analysis

*S. aureus* MVs, cell lysates, or serial dilutions of filter-sterilized biofilm homogenates before and after ultracentrifugation were heated at 95°C for 30 min to denature proteins. Samples were applied to nitrocellulose membranes using a 96-well Bio-dot apparatus. The membranes were blocked with 5% skim milk containing rabbit IgG and incubated overnight at 4°C with one of the following primary antibodies: human PNAG monoclonal antibodies (2 μg/mL), CP8 mouse monoclonal antibodies (5A6; 0.5 μg/ml), or LTA mouse monoclonal antibodies (0.5 μg/mL, IBT Bioservices). After washing, membranes were incubated with the appropriate HRP-conjugated secondary antibodies (1:5000) and developed using a chemiluminescent HRP substrate.

### Confocal microscopy

Biofilms grown in 8-well chambered coverglass or on glass slides were gently washed 2-3 times with sterile PBS and fixed for 30 min with 4% formaldehyde at room temperature. The biofilms were then rinsed with PBS and stained for 30 min with 5 μM DNA dye SYTO-9 (Thermo Fisher). Imaging was performed with Nikon NIE upright confocal microscope. Z-stack images were acquired at 1.0 µm intervals through the biofilm depth. The thicknesses (μm) and biovolumes (μm^3^ μm^−2^) of the biofilms were measured using the COMSTAT2 software (www.comstat.dk).

### eDNA measurement

To compare the amount of eDNA associated with MVs produced from drip-flow biofilms and planktonic cultures, DNA was extracted from equal amount (5 μg) of MVs using a Monarch® Spin gDNA Extraction Kit (NEB, MA) according to manufacturer’s instruction and quantified with a NanoDrop Spectrophotometer (Invitrogen). To determine the proportion of eDNA associated with MVs versus that freely present in the biofilm matrix, equal volumes (30 μL) of filter-sterilized biofilm homogenates collected before (fraction S1) and after MV depletion (fraction S2) by ultracentrifugation were loaded onto a 1% agarose gel and stained with SYBR Safe DNA gel stain. DNA signal intensity was quantified using ImageJ (National Institutes of Health, Bethesda, MD, USA).

Alternatively, 100 μL of serially diluted S1 and S2 fractions or PBS control were mixed with an equal volume of 4 μM SYTOX Green DNA dye and incubated for 15 min at room temperature. Fluorescence intensity was measured using a plate reader (Infinite 200 Pro, Tecan) with excitation and emission wavelengths of 465 nm and 510 nm, respectively. Background fluorescence from the PBS control was subtracted from each sample reading.

## Data availability

The mass spectrometry proteomics data were deposited to the ProteomeXchange Consortium via the PRIDE partner repository with the data set identifier PXD075156.

## Acknowledgement

We thank Dr. Jinpeng Liu for bioinformatic support through the Center for Computational Sciences at the University of Kentucky. We thank Maria Ericsson from Electron Microscopy Core at Harvard Medical School for help with TEM imaging of ultrathin sections. We also thank Gerald Pier and Angelika Gründling for generously providing human PNAG monoclonal antibodies and *S. aureus* LAC Δ*ltaS* suppressor strains, respectively. We are grateful to Dr. Jean C. Lee for valuable advice on the manuscript. This work was supported by the National Institute of Allergy and Infectious Diseases of the NIH under award no. R21AI180326 (X.W.). X.W. conceived the project. J.L., E.G., and X.W. designed the experiments. J.L., X.W., M.F., and E. N. performed experiments. J.L., X.W., and E.G. analyzed the data. X.W. and J.L. drafted the manuscript. All authors reviewed and edited the manuscript.

